# Rapid evolution of ecological sexual dimorphism driven by resource competition

**DOI:** 10.1101/2022.05.09.491145

**Authors:** Stephen P. De Lisle

## Abstract

Sex differences in ecologically-important traits are common in animals and plants, and prompted Darwin to first propose an ecological cause of sexual dimorphism. Despite theoretical plausibility and Darwin’s original notion, a role for ecological resource competition in the evolution of sexual dimorphism has never been directly demonstrated and remains controversial. I used experimental evolution in *Drosophila melanogaster* to test the hypothesis that resource competition can drive the evolution of sex differences in diet. Following just three generations of adaptation, offspring from flies evolved in low-resource, high-competition environments show elevated sexual dimorphism in diet preference compared to both the ancestor and populations evolved on high resource availability. These results provide the first real-time direct evidence for evolution of sexual dimorphism driven by an ecological cause.

**One sentence summary:** Sex differences in fly diet evolved rapidly under elevated competition, demonstrating ecological cause of sex differences.

## Main Text

Trait differences between the sexes, or sexual dimorphisms, are a striking source of diversity in sexually-reproducing organisms (*1, 2*). Although these sexual dimorphisms often reflect traits related to mating, in many cases males and females have also diverged in traits related to ecological niche, such as body size, diet content, habitat use, and feeding morphology (*3*). This diversity of sexual dimorphism prompted Darwin (*2*) to propose, in addition to sexual selection, the possibility of an ecological cause of sexual dimorphism (*4*). In modern terms this “ecological cause” has been framed as a potential role for resource competition in driving ecological character displacement between the sexes (*5*). Mathematical models suggest character displacement between the sexes can readily evolve (*5–8*), and a number of studies in birds (*9*), salamanders (*10*), lizards (*11–13*) and other systems (*14*) suggest that competition may play a role in the evolution of sexual dimorphism. However, unlike other causes of the evolution of sexual dimorphism such as sexual selection, Darwin’s ecological cause of sexual dimorphism remains controversial and lacks unambiguous evidence (*14*).

Laboratory experimental evolution provides a powerful approach for testing the hypothesis that ecological conditions may drive the evolution of sexual dimorphism. In *Drosophila* flies (*15–18*) and other insects (*19*), male and female fitness are typically maximized under different ratios of protein and carbohydrate (*20*), with female fecundity maximized under relatively high protein diets and male survival and mating success maximized under relatively high carbohydrate. However, in multiple *Drosophila melanogaster* populations (e.g., (*15*) and the ancestral population of this study, Figure 1), standing variation in adult male and female diet preference remain largely overlapping with non-significant sexual dimorphism, implying that ecological conditions may determine the extent to which sexual dimorphism in diet preference evolves. Resource competition at the larval stage has been shown to be an important driver of adaptation to resource use in *D. melanogaster* (*21*). Thus, D*. melanogaster* makes an ideal model system to experimentally test the hypothesis that resource competition at the adult stage can drive character displacement between the sexes, with the *a priori* prediction that males and females will alternatively evolve elevated carbohydrate and protein source consumption, respectively, under elevated resource competition.

**Fig. 1.**
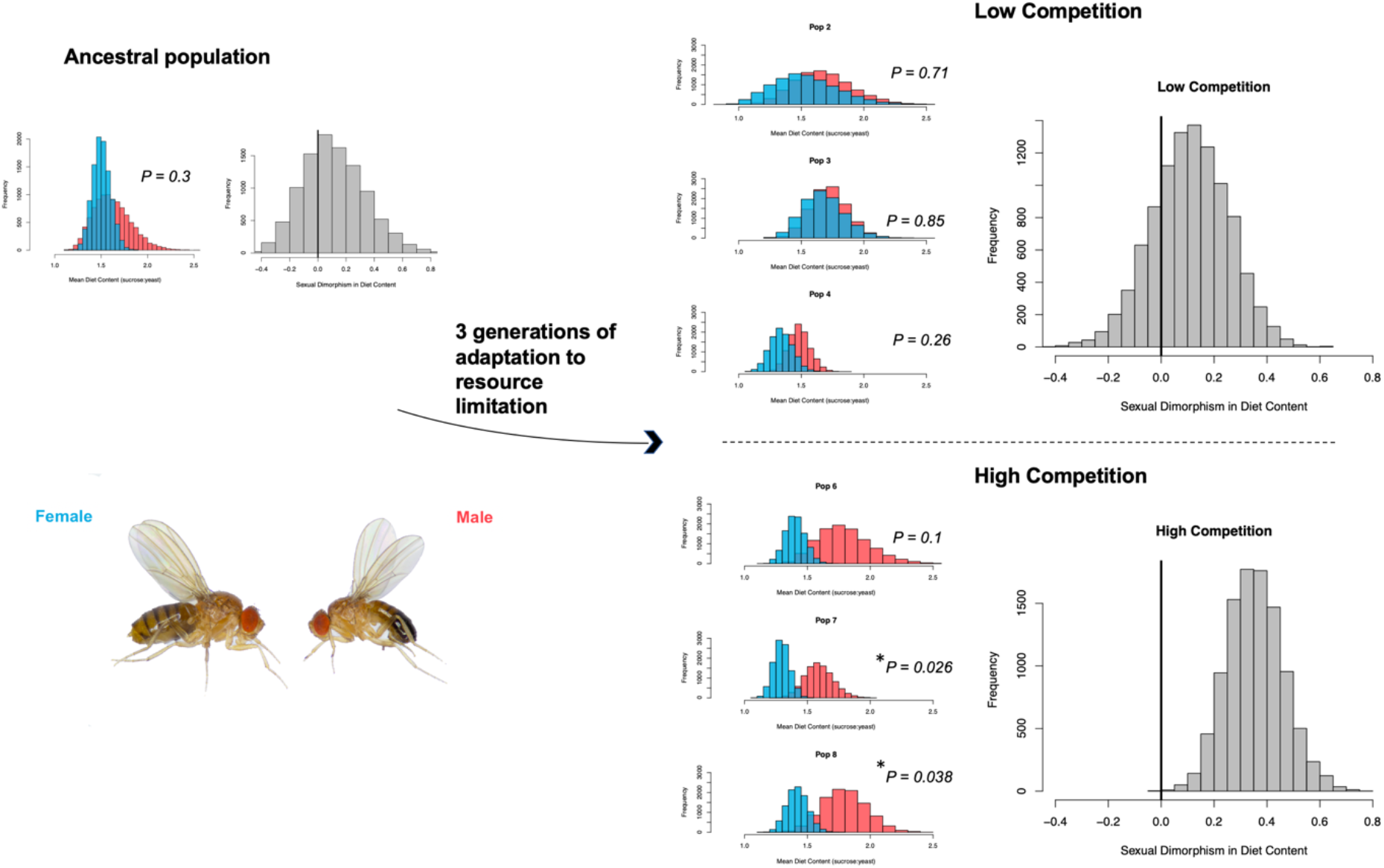
Sexual dimorphism in diet content evolved from a monomorphic ancestor in three generations. Sampling distributions of male and female mean sucrose:yeast consumption ratios show large overlap between male and female diet in the ancestral population (left panel), and sexual dimorphism was not statistically significant (sampling distribution of the difference in male and female mean shown). After 3 generations of adaptation to high competition (low resource availability), 2 populations showed significant sexual dimorphism and the third replicate was trending towards significant dimorphism. On average, populations adapted to high competition showed significant sexual dimorphism, with males consuming higher sucrose:yeast ratios than females (right panel, bottom). Sexual dimorphism was not significantly different from zero in populations adapted to low competition, high resource availability (upper right). P-values are from linear models with sex as a main effect and sucrose:yeast ratio as the response, fit separately for each population. Sampling distributions were calculated by bootstrapping, stratified by population for pooled sampling distributions of the difference.

I evolved 6 replicate *D. melanogaster* (LHm background, *22*) populations under conditions of elevated adult resource competition. By housing flies in vials where larval food was covered with a thin layer of agar (readily penetrated by hatched larvae but not by adult flies; see Supplementary Material), I was able to manipulate adult liquid diet independently of larval diet using microcapillary tubes; adults in half of the populations received 5 μL of sucrose (carbohydrate source) solution and 5 μL of yeast (protein source) solution, while the other three populations evolved under conditions of 30 μL of sucrose solution and 30 μL of yeast solution at constant adult density. These treatments thus correspond to low resource availability, high competition, and high resource availability, low competition respectively. These conditions were replicated each generation, with standard larval food constant across the two treatments.

After just three generations, offspring from all populations evolving under high competition (low resource availability) showed elevated sexual dimorphism in sucrose:yeast consumption, with males showing increased sucrose:yeast ratios compared to females (Figure 1) consistent with predictions. This sexual dimorphism was statistically significant or trending towards significance in all three replicate populations (Figure 1), and the mean estimate from all high competition populations indicate that on average sexual dimorphism in diet content was highly significant (linear mixed effects model, *F_1, 236_* =11, *P* = 0.001). Populations evolving under low competition (high resource availability) did not show significant sexual dimorphism (linear mixed effects model, *F_1,195_* = 0.76, *P* = 0.38, Figure 1); this treatment effect on sexual dimorphism was statistically significant (linear model *F_1, 4_* = 25.4, *P* = 0.007).

Mean fitness of all six populations increased an order of magnitude during the course of the experiment, indicating evolution was adaptive (Figure 2A). Survival during 48 hours of food deprivation was higher for offspring from high competition, low food environments than for flies evolving in high food conditions (generalized linear model *Z* = 4.5, *P* < 0.0001, Figure 2B), indicating that high competition populations evolved adaptation to resource restriction. These patterns of elevated resource acquisition and survival under resource restriction are consistent with expectations from functional relationships between resource acquisition and starvation resistance in *D. melanogaster* (*18*).

**Fig. 2.**
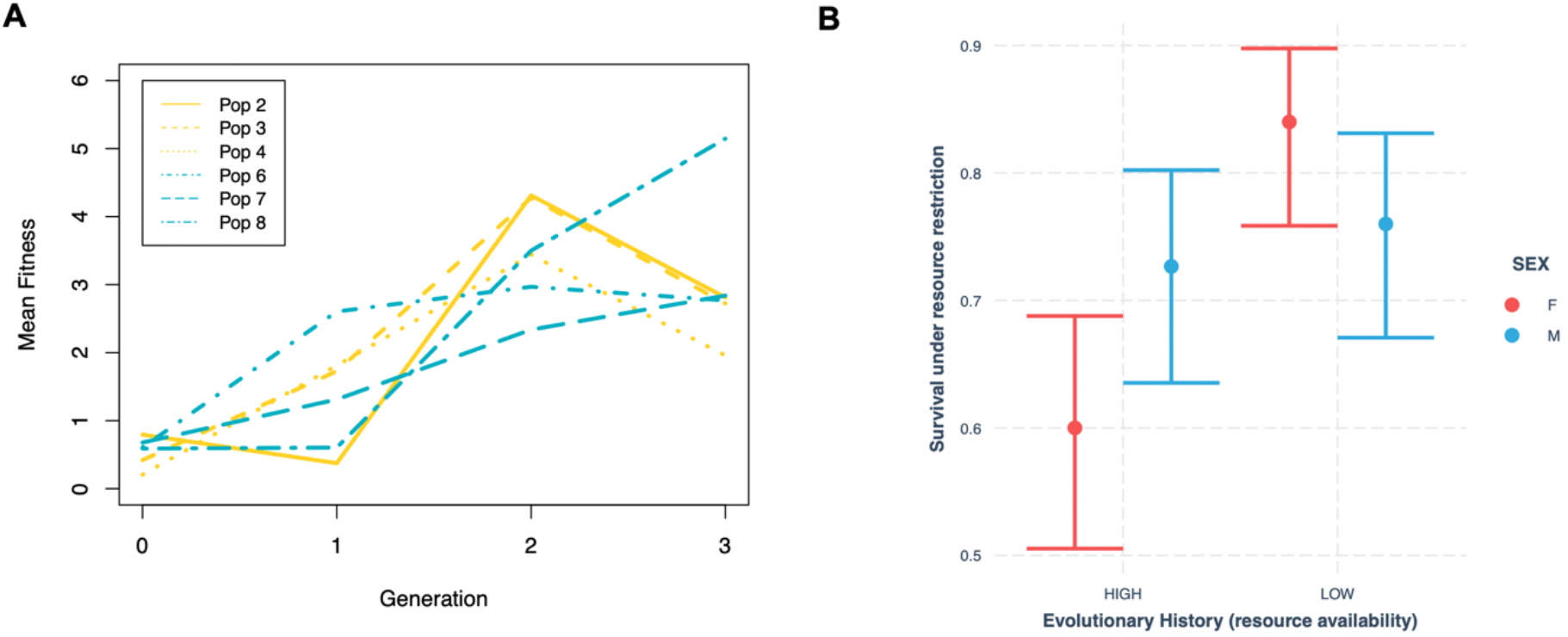
Populations adapted to environmental conditions over the course of the experiment. Panel **A** shows mean fitness (calculated from recruitment to the next generation) of each population from the onset of the experiment; mean fitness increased for all six populations over the course of the experiment. Panel **B** shows survival (estimates and 95% CIs from a binomial glm) during 48 hours of resource restriction in offspring from generation 3 flies. Populations adapted to low resource availability (high competition) showed elevated survival under resource restriction compared to flies evolved under high resource availability.

An analysis of bivariate diet content (*23*) revealed that populations evolved under resource restriction evolved elevated levels of resource consumption compared to high resource populations (bivariate mixed effects model, posterior mean treatment effect = 25.9 (3.7 – 47 95% CI) micrograms, *pMCMC* = 0.034; Figure 3A), indicating adaptation to elevated competition. Males evolved elevated sucrose intake while females evolved elevated yeast intake (bivariate mixed effects model, posterior mean trait*sex = −48 (−84 – −13 95% CI) micrograms, *pMCMC* = 0.014; Figure 3A), and evolutionary rates were not significantly different between the sexes (sucrose: linear model *F_1,10_* = .96, *P* = 0.34, yeast: *F_1, 10_* = 1.8, *P* = 0.21, Figure 3B), indicating that evolution of sexual dimorphism in diet was not driven by divergence in one sex, but rather by divergence in both male and female diet.

**Fig. 3.**
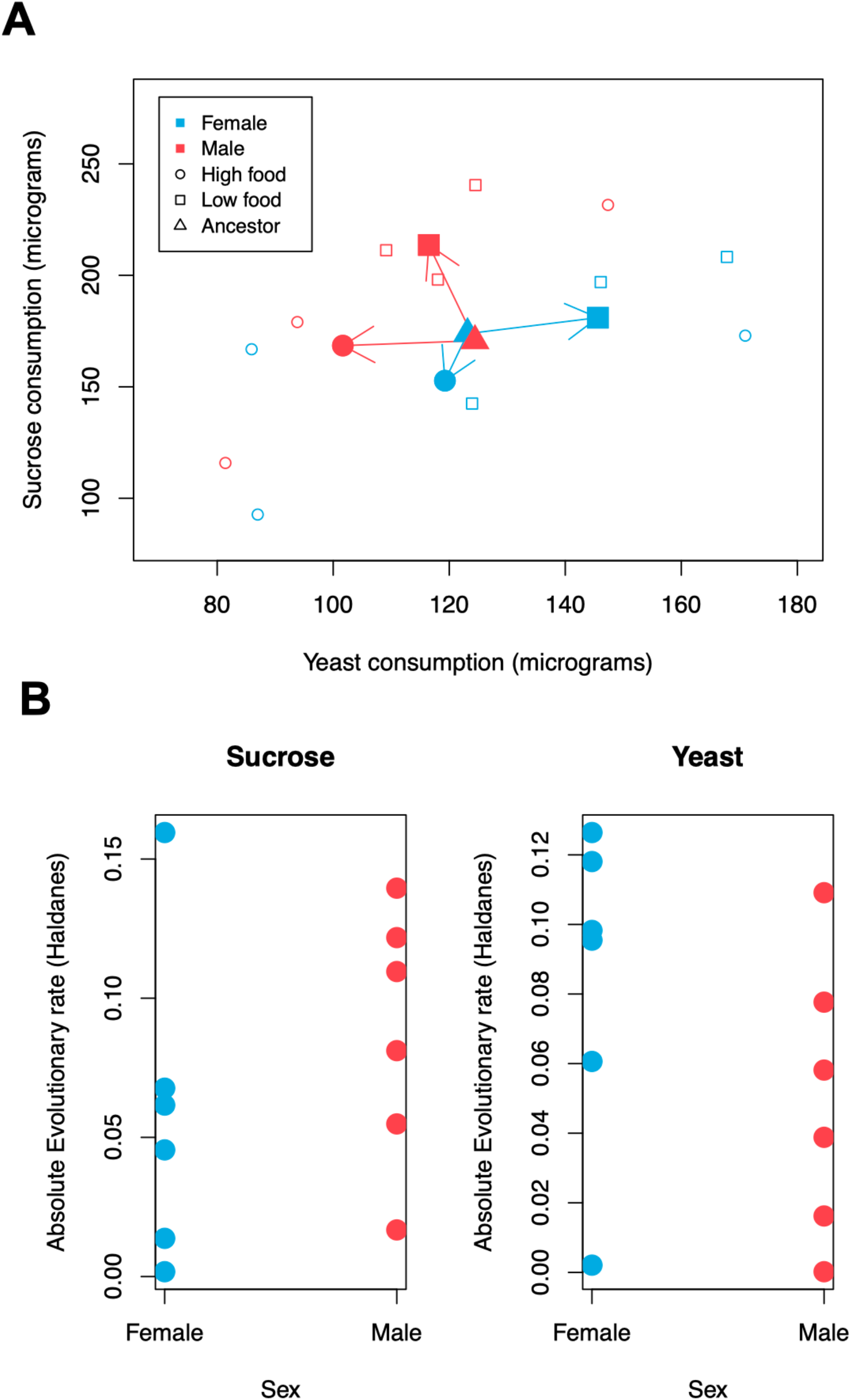
Evolution of sexual dimorphism occurred via divergence in both male and female diet. Panel **A** shows mean sucrose and yeast consumption for offspring from males and females from each population (small open markers) after 3 generations of adaptation, as well as the treatment means (large filled markers). Arrows connect male and female ancestral means to the corresponding means for populations adapted to high and low resource availability. Panel **B** shows the absolute value of the evolutionary rate in Haldanes for each sex for each population, measured as 1/3 of the difference between generation 3 mean diet content and the ancestral mean over the ancestral standard deviation.

Theory suggests that competition can drive sexual dimorphism in resource use most readily when the sexes interact in small demes with reliably-distributed resources (*6*). The experiment reported here is consistent with this expectation in that the design entailed interactions occurring in demes (vials) of 8 males and 8 females, with consistent spatial distribution of resources across vials and generations within a population. Although male-specific displacement is expected under conditions of monogamy (*6*), *D. melagnogaster* are not monogamous (*24*), and the finding of displacement in both sexes is thus consistent with our experimental design and predictions of character displacement theory. The predictable direction of the evolution sexual dimorphism under high competition (elevated male sucrose intake, elevated female yeast intake) in this experiment indicate that resource competition acts along with nutritional requirements related to sex specific reproductive strategies to influence male and female diet evolution (*14*).

This experiment demonstrates that an environment of elevated resource competition can drive the rapid evolution of ecological sexual dimorphism from a non-dimorphic or minimally-dimorphic ancestor, as one component of adaptation to elevated adult resource competition. Although these results are under laboratory conditions in a model organism, evolutionary rates of sucrose (0.072 Haldanes, average) and yeast (0.066 Haldanes, average) consumption in this experiment (Figure 3B) are well within the range observed in contemporary studies of microevolution in the wild (*25*). Combined with the prevalence of ecologically-important sex differences from a diverse range of taxa (*3, 9, 14*), evidence from natural systems such as competition-driven divergent selection (*10*) and geographic patterns consistent with character displacement between the sexes (*11, 13*), the experiment presented here provides a proof-of-concept that demonstrates that Darwin’s idea of an ecological cause of sexual dimorphism may be an important and general contributing cause to the diversity of sexual dimorphism observed in nature.

## Aknowledgements

I thank K. Lund-Hansen, A. Singh, E. Svensson, J. Abbott, N. Feiner, R. Prantner, and M. Frieberg for sharing knowledge and/or resources.

## Data and Materials Availability

All data is available in the manuscript or the supplementary materials.

## Supplementary Materials

### Materials and methods

#### Fly rearing and experimental design

Experimental populations were derived from a LH_m_ stock population of *D. melanogaster*. This population is from a lab-adapted population originally collected in California in 1991 from 400 females (*22, 26*), and obtained by me from Jessica Abbott’s lab group at Lund University, where their population has been maintained for over 500 generations. This population was maintained under conditions of 12:12 light, approximately 25 C, 60% humidity and fed standard fly medium consisting of cornmeal, yeast and molasses. As a standard component of the rearing protocol, yeast is typically added for consumption by newly eclosed adults to stimulate fecundity; thus, the population is adapted to high resource availability at the adult stage. One generation prior to the start of the experiment, diet preference was assayed by placing 50 males and 50 females in individual fly vials containing 5mL of agar solution (to prevent desiccation) along with 2 5μL microcapillary tubes, one containing 90g/L sucrose solution and the other containing 90g/L yeast extract solution (ultra pure bacteriological grade, VWR J850). Flies were removed after 24 hours and the length of fluid remaining in the microcapillary tubes was measured to the nearest hundredth of a millimeter with digital calipers.

Eight populations were established with 7 28mm wide fly vials each, with 8 males and 8 females per vial, for a total of 112 flies per population (Table S1). Vials contained standard fly food (approx. 15 mL) to which the LH_m_ population is highly adapted, capped with a thin layer (approx.. 3 mL) of agar solution to block access of adult flies to nutrition present in the larval food source. Larvae can readily penetrate this agar cap; a pilot experiment showed that effects of the larval cap on vial reproductive output were minimal (mean eclosed from agar-capped vials following oviposition by 5 females: 117, regular vials: 130, *F_1,8_* = 1.66, *P* = 0.23). Microcapillary tubes were added to the vials for 48hours to provide a source of carbohydrate and protein for adult flies. After 48 hours, adults and microcapillary tubes were removed from the vials and resulting offspring allowed to develop. This process was then repeated in the next generation; at each generation, newly eclosed flies were removed from vials, pooled by population, counted, sexed and allocated randomly under light C0_2_ anesthesia to new agar-capped vials. Density was kept constant at 16 flies per vial, and population size was allowed to expand by expanding the number of vials in a population up to the founding size of 7 vials. Four high resource availability populations received 30 μL of sucrose solution and 30 μL of yeast solution (both 90g / L) in a total of 6 10 μL microcapillary tubes. Four low resource availability populations received a single 5 μL microcapillary tube each of sucrose solution and yeast solution. One population from each treatment group failed to produce enough flies in the first generation to yield a single vial at the standard density and so were treated as extinction events.

After 3 generations of selection, experimental flies were placed in regular food vials after removal from their agar-capped experimental vials to produce offspring reared in a common environment for diet assay. These offspring were collected upon eclosion and placed individually in vials with 5 mL agar solution to prevent desiccation. Flies were held for 48 hours after which 2 5μL microcapillary tubes, one containing 90g/L sucrose solution and the other containing 90g/L yeast extract solution, were added. 20 additional vials, lacking flies, were dispersed throughout the environmental chamber and received microcapillary tubes in order to control for effects of evaporative loss. Flies were removed after 24 hours and the length of fluid remaining in the microcapillary tubes was measured to the nearest hundredth of a millimeter with digital calipers.

A predetermined stopping point of three generations was determined in advance; offspring counts after a 4^th^ generation of treatment exposure were conducted to obtain recruitment data needed for a mean fitness measure for generation 3, it was not possible to further assay diet due to logistical constraints.

#### Controlling for Evaporation

Evaporative loss was high in this experiment; past experiments using liquid media consumption from microcapillary tubes to measure diet in *D. melanogaster* have used very high relative humidity (>80%, (*15, 16*)), which was not desirable in this design because it would conflate the diet change of the experimental manipulation with a significant increase in relative humidity, since the ancestral population is highly adapted to 60% relative humidity. I took two approaches to correct for evaporative loss. First, in the analysis of sexual dimorphism in the ratio of sucrose:yeast consumption, I corrected raw ratios by the ratio of evaporative loss from control vials, as (sucrose_raw_:yeast_raw_ – sucrose_evaporation_:yeast_evaporation_) + 1; equivalent conclusions were obtained using the raw values. For the analysis of bivariate sucrose and yeast consumption, in which negative values for some small number of individuals (arising due to measurement error) do not pose the conceptual or mathematical problem that they do for ratios, I subtracted sucrose_evaporation_ and yeast_evaporation_ from the respective raw values. Corrected values for all populations were significantly greater than zero (all *P* ≤ 0.00017), indicating that this measure of fluid loss is indeed capturing consumption of individual flies; equivalent general conclusions were obtained in analysis of the raw length of fluid remaining.

#### Statistical Analysis

I used separate linear models for each population to assess the effect of sex on the ratio of sucrose:yeast consumption on a population by population level. To assess average effects by treatment, I used two separate mixed effects models with sex as a main effect and population as a random effect. To generate treatment-wide sampling distributions for sexual dimorphism in sucrose:yeast (Figure 1), I bootstrapped each treatment group using stratified (by population) random sampling with replacement (10,000X). To estimate treatment effects on sexual dimorphism while avoiding fitting a model with an interaction term, for which power would be low with just 6 populations, I first bootstrapped (10,000X) the data to obtain mean and variance estimates of male sucrose:yeast – female sucrose:yeast, and then fit a mixed model with this population mean measure of sexual dimorphism as the response, treatment as the main effect, and weighted by the inverse variance of population sexual dimorphism. To assess mortality effects, I fit a binomial glm with each populations’ survival and mortality as the response. To analyze bivariate response in sucrose and yeast consumption, I fit a series of Bayesian multiresponse mixed effects models with varying degrees of complexity; a model with just trait-specific intercepts, a model with treatment effect, sex effect, treatment*trait+sex, and and sex*trait + treatment. All models contained a random trait-specific intercept among populations and an unstructured residual covariance matrix; default priors were used. Of these models the model with sex*trait interaction and main effect of treatment fit best (ΔDIC = 2.7). All statistical analyses were performed in R; linear models were fit using lm(), mixed effects models using lme(), and multiresponse mixed effects models were fit using MCMCglmm. Complete R script is provided as supporting material.

**Table S1.** Summary of Population Sizes at each Generation.

| **Generation 0** |  |  |
| --- | --- | --- |
| **population** | **N** | **vials** |
| 1 | 112 | 7 |
| 2 | 112 | 7 |
| 3 | 112 | 7 |
| 4 | 112 | 7 |
| 5 | 112 | 7 |
| 6 | 112 | 7 |
| 7 | 112 | 7 |
| 8 | 112 | 7 |

| **Generation 1** |  |  |
| --- | --- | --- |
| **population** | **N**<br>**eclosed** | **vials for next**<br>**generation** |
| 1 | 6 | 0 |
| 2 | 89 | 5 |
| 3 | 47 | 3 |
| 4 | 23 | 1 |
| 5 | 0 | 0 |
| 6 | 68 | 3 |
| 7 | 76 | 4 |
| 8 | 66 | 3 |

| **Generation 2** |  |  |
| --- | --- | --- |
| **population** | **N**<br>**eclosed** | **vials for next**<br>**generation** |
| 2 | 30 | 1 |
| 3 | 83 | 5 |
| 4 | 29 | 1 |
| 6 | 125 | 6 |
| 7 | 84 | 5 |
| 8 | 29 | 1 |

| **Generation 3** |
| --- |

| **population** | **N**<br>**eclosed** | **vials for next**<br>**generation** |
| --- | --- | --- |
| 2 | 69 | 4 |
| 3 | 342 | 7 |
| 4 | 55 | 3 |
| 6 | 285 | 7 |
| 7 | 187 | 7 |
| 8 | 56 | 3 |

**Generation 4**
| **population** | **N**<br>**eclosed** | **vials for next**<br>**generation** |
| --- | --- | --- |
| 2 | 180 | NA |
| 3 | 305 | NA |
| 4 | 94 | NA |
| 6 | 310 | NA |
| 7 | 318 | NA |
| 8 | 247 | NA |

